# Psilocybin causes sex, time, and dose dependent alterations in brain signaling pathways

**DOI:** 10.1101/2024.12.16.628764

**Authors:** J. Hudson Barnett, Kennedi T. Todd, Joseph Benetatos, Benjamin E. Rabichow, Katelin A. Gibson, Kimberly C. Olney, John D. Fryer

## Abstract

Psilocybin is a psychedelic tryptamine that has emerged as a potential candidate for the treatment of a variety of conditions, including treatment resistant depression and post-traumatic stress disorder. Clinical trials which have assessed the efficacy of psilocybin for these conditions report a rapid and sustained improvement in patient- and clinician-rated depression scores. The established mechanism of action for psychedelics such as psilocybin is agonism of the serotonin 2A receptor (5HT_2A_R), however, the downstream events that mediate their therapeutic effects remain uncertain. As high doses of psychedelics are known to induce strong perceptual alterations, an additional outstanding question is whether subperceptual doses induce similar molecular effects as psychoactive dosages. Here, we report the first analysis of dose- and sex-dependent transcriptional changes in forebrains of female and male mice at 3 timepoints (8 hours, 24 hours, and 7 days) following a single administration of psilocybin at low (0.25 mg/kg) or high (1 mg/kg) doses. Grouped analysis of both sexes reveals dose- and time-dependent transcriptomic alterations. We report more rapid transcriptional changes and attenuation of such changes in females following a single low-dose relative to males treated identically. Females also responded more robustly to high-dose administration relative to males at 8 and 24 hours, with signal attenuation in both sexes by 7 days. A notable observation was the persistent transcriptional effect of low-dose psilocybin at 7 days, which outlasted high-dose changes, and which suggests that low doses may have prolonged biological effects. A myriad of pathways were altered depending on sex and timepoint, but common features included functions related to neuronal differentiation, neurogenesis, and changes in receptor signaling. These data reveal dose- and sex-dependent molecular effects of psilocybin and support previous studies demonstrating its effect on dendritogenesis. Given ongoing clinical interest in psilocybin for treating mental health disorders, our results suggest that these sexually divergent changes should be considered when weighing treatment strategies. Additional consideration should be given to temporal effects of low vs high dosages on gene transcription, especially when timing psilocybin with adjuvant cognitive behavioral therapy.

## Introduction

Psilocybin – a psychedelic tryptamine prodrug found in over 200 species of fungi – is currently under phase 3 clinical investigation for major depressive disorder (NCT06308653), treatment resistant depression (TRD) (NCT05624268), and in phase 2 trials for post-traumatic stress disorder (PTSD) (NCT05312151). The clinical efficacy of psilocybin was explored extensively in the 1960s following the discovery of its profound effects on perception, mood, and cognition^1^. This naturally occurring compound undergoes hepatic dephosphorylation to yield psilocin, the active metabolite with high affinity for serotonin (5-HT) receptors^2^.

Serotonin is a central neurotransmitter involved in the regulation of emotion, mood, appetite, social interactions, and sleep^3,4^. Understanding serotonergic neurotransmission in the brain is particularly important within the context of disordered psychiatric systems. Excluding ionotropic 5-HT_3_, the 5-HT receptor family are transmembrane G protein-coupled receptors (GPCRs) that mediate signaling cascades in response to hormones, neurotransmitters, and environmental stimuli^5^. Binding of 5-HT receptors by psilocin and serotonin activate intracellular signaling cascades via biased agonism of GPCRs, resulting in downstream molecular and behavioral outcomes^6–8^.

Modulation of serotonin signaling has been an important treatment strategy for psychiatric disorders since the FDA approved fluoxetine, a selective serotonin reuptake inhibitor (SSRI)^9^. This class of drugs is the first-line choice for pharmacologic management of PTSD, major depressive, generalized anxiety, obsessive compulsive, and eating disorders^10^. Their efficacy has been attributed to increased availability of synaptic serotonin, prolonging interactions within its receptor system^11^. The action of psilocin within the 5-HT receptor system has been revealed through a series of foundational studies, establishing the 5-HT_2A_ receptor as the primary mediator of psychedelic effects^6,8,12,13^. However, the mechanistic basis of how psilocybin produces an antidepressant effect within hours and sustains it for >1 month is still open to investigation. For instance, a recent study revealed that the therapeutic properties of both SSRIs and psilocybin may depend on allosteric modulation of the TrkB receptor^14^. This discovery reinforces the need to further characterize the molecular effects of emerging therapies such as psilocybin, which may have antidepressant properties due to interactions beyond the 5-HT receptor system.

Preclinical investigations of psilocybin therapy have employed a high dose (1 mg/kg or more) at a single timepoint to assess the neuroplastic mechanisms and behavioral effects^14–17^. Data from human clinical trials indicate both acute and chronic effects following a single dose of psilocybin^18–20^. In the present study, we expand the purview of psilocybin’s known sex-, dose-, and time-dependent effects on the mammalian brain by assessing transcriptional alterations in murine forebrains following single-dose-intraperitoneal treatments.

## Methods

### Resource Availability

#### Lead Contact

Requests for further information and resources should be directed to and will be fulfilled by the lead contact, John D. Fryer

#### Materials availability

This study did not generate new reagents or materials

#### Data and code availability

The data for these findings and the code to analyze these findings is available at https://github.com/fryerlab/psilocybin. Upon manuscript acceptance, raw data will be available via GEO and SRA.

### Method details

#### Psilocybin

A DEA schedule 1 license was procured and used to obtain psilocybin (cat #7437-001) from the National Institute on Drug Abuse, Drug Supply Program (Bethesda, MD, USA). The chemical composition and purity of psilocybin was verified using high pressure liquid chromatography. Psilocybin was suspended in 0.9% saline and administered via intraperitoneal injection.

#### Experimental design

Female and male C57BL/6J adult mice (The Jackson Laboratory, strain # 000664) at 10 weeks of age were used. Animal housing and experimental protocols were conducted within the Mayo Clinic Arizona under an approved IACUC protocol (# A00007629-24). Animals were moved from the vivarium and allowed to habituate in the procedure room for 30 minutes. Animals received a 0.2 mL i.p. injection of low (0.25 mg/kg) or high (1 mg/kg) dose psilocybin or vehicle-control (0.9% saline) using N=5 mice/sex/group and were then returned to their home cages while awaiting sacrifice at three pre-determined timepoints. At designated sacrifice timepoints, mice received a lethal dose of euthasol (0.03 mL) and anesthetic depth was assessed via toe pinch. Once animals were no longer responsive, transcardial perfusion was performed with 15 mL of phosphate buffered saline (PBS) with 5 mM EDTA. The hemiforebrain was collected and flash frozen on dry ice prior to being transferred to long term storage at -80 °C.

#### Bulk RNA-seq and statistical analysis

A total of 0.5 grams of forebrain tissue was used to prepare total brain RNA with the Qiagen RNeasy Mini Kit (cat # 74104). Samples were QC’d on the Agilent Tapestation, and bulk RNAseq libraries prepared with TruSeq stranded mRNA (cat # 20020594) and sequenced to a depth of ∼50M paired-end reads on an Illumina NovaSeq S4 instrument at the Mayo Clinic Genome Sequencing Core. FASTQ files were trimmed for adapter content and polyA/polyG artifacts using BBDuk^21^. FastQC and MultiQC were used to assess read quality before and after trimming^22,23^. Trimmed reads were aligned to reference genome GRCm38 version M23 (Ensembl 98) using STAR^24^. Gene features were counted using featureCounts from the Subread package^25^. A counts matrix of gene rows by sample columns was created and read into R for analysis. Mitochondrial genes were removed from the analysis and only protein coding genes were retained. Lowly expressed genes were removed from the analysis by keeping only genes that had a count greater than or equal to one count per million (CPM) in all samples within at least one group (defined by time point/dose or time point/dose/sex in sex-specific analyses). The edgeR package converted library sizes into effective library sizes using the Trimmed Mean of M-values (TMM)^26^. QC was done before and after filtering lowly expressed genes and TMM normalization. For analyses including both sexes, a model with group (defined by time point and dose) and log2(*Hbb-bs* CPM) was used. *Hbb-bs* was added as a covariate to account for subtle differences in perfusion. For sex-specific analyses, a model with group (defined by time point, dose, and sex) and log2(*Hbb-bs* CPM) was used. Differential expression was performed using the limma/voom pipeline^27,28^. P-values were adjusted for multiple tests using the Benjamin-Hochberg correction.

#### Metascape gene-set enrichment analysis

Metascape gene annotation analysis and enrichment^29^ was used to highlight pathways that undergo differential regulation in psilocybin treated versus vehicle-control animals. Metascape Z-scores (Z-standard deviation away from expected counts) are calculated by comparing the overlap of an input set of differentially expressed genes to specific gene ontology term to assess deviation from expected overlap due to random chance. A larger Z-score (negative or positive) denotes greater deviation from the expected distribution.

#### Weighted Gene Co-Expression Network Analysis

A Weighted Gene Co-Expression Network Analysis (WGCNA) was performed using the WGCNA R package^30^ to identify clusters of co-expressed genes associated with key phenotypic traits. The soft-thresholding power was chosen based on the scale free topology model fit (R2) just above 0.8.

## Results

### Low and high dose psilocybin produce distinct effects on the transcriptome

To assess the acute effects of psilocybin at different doses using RNA sequencing (RNA-seq) of forebrain tissue, we first analyzed male and female mice grouped together to determine differentially expressed genes (DEGs) following a low or high dose of intraperitoneal (i.p.) psilocybin, along with saline controls, using N=5 mice/sex/group. Assessing the male and female animals together, we observed transcriptional changes at both doses compared to vehicle (0.9% saline) at 8 hours following psilocybin administration. The effect of low dose psilocybin at 8-hours was minimal, with more numerous changes resulting from high dose psilocybin. Animals in the low dose group showed only 2 significant DEGs compared to vehicle (Supp Fig 1A). However, mice receiving high dose psilocybin showed dozens of changes, with slightly more downregulated than upregulated DEGs (Supp Fig 1B). Pathways impacted by the downregulated transcripts included small molecule transport, nucleosome assembly, and regulation of neurogenesis (Supp Fig 1C, D). Given the smaller number of upregulated DEGs, only a single pathway was significantly altered: multicellular organismal-level homeostasis (Supp Fig 1C).

**Figure 1.**
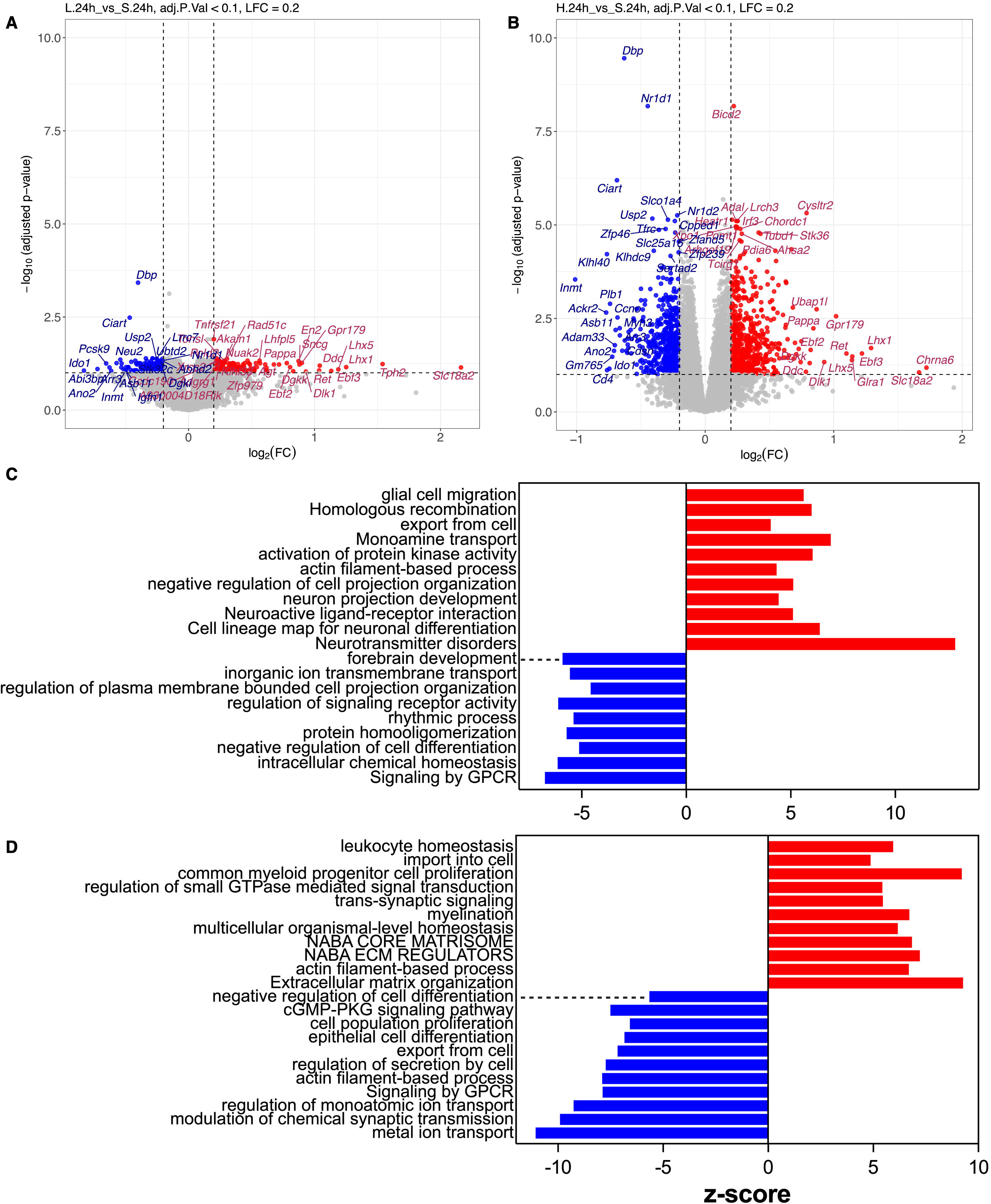
High and low dose psilocybin produce distinct and overlapping transcriptomic changes at 24-hours. (A,B) Bulk RNA-seq volcano plots. Downregulated (blue) and upregulated (red) CNS transcriptional changes 24-hours following (A) 0.25 mg/kg i.p. psilocybin compared to vehicle (0.9% saline) and (B) 1 mg/kg i.p. psilocybin compared to vehicle. Log2 fold-change (FC) denotes magnitude of transcript differential expression on the x-axis. -log10(adjusted p-value) shows FDRq corrected p-values. (C,D) Gene Ontology (GO) analysis characterizes the most significantly down- and upregulated pathways 24-hours following (C) low and (D) high dose psilocybin. Z-score denotes Z-standard deviation away from expected counts on the x-axis, and GO pathways on the y-axis.

To assess how the effects of psilocybin evolved over time, we measured transcriptional changes 24 hours following low and high dose i.p. psilocybin. At 24-hours, low dose psilocybin altered gene expression but with far fewer DEGs compared to high dose (Fig 1A-B). Significant overlap was seen between low and high doses, with 115 shared upregulated and 110 shared downregulated transcripts (Supp Fig 2), which included transcripts with the largest fold change. Circadian genes *Ciart*, *Dbp*, and *Nr1d1* remained downregulated at the 24-hour timepoint indicating that alterations to sleep cycles may be a shared feature of low and high doses psilocybin (Fig 1A-B). Shared upregulation of neuronal transcription factors *Lhx1*, *Lhx5*, *Ebf3*, and transmembrane monoamine transporter *Slc18a2* highlight the overlapping effects of low and high dose psilocybin on inducing pathways of neuronal differentiation and monoamine transport (Fig 1A-B). While low and high dose psilocybin induced overlapping down- and upregulated DEGs at 24-hours, each dose was also associated with distinct, nonoverlapping patterns of gene expression (Supp Fig 2). Distinct effects of low dose psilocybin that were not present in high dose animals included 105 down- and upregulated transcripts (Supp Fig 2) involved in neuron projection development, synapse organization, and cytosolic calcium regulation (Fig 1C). High dose psilocybin induced far more distinct changes at 24 hours compared to low dose, with 903 down- and upregulated transcripts (Supp Fig 2) involved in cell proliferation, extracellular matrix (ECM) remodeling, and homeostatic processes related to Forkhead box O (FOXO) transcription (Fig 1D). These findings suggest that lower doses of psilocybin have distinct molecular effects while simultaneously altering a subset of the genes impacted by high doses.

**Figure 2.**
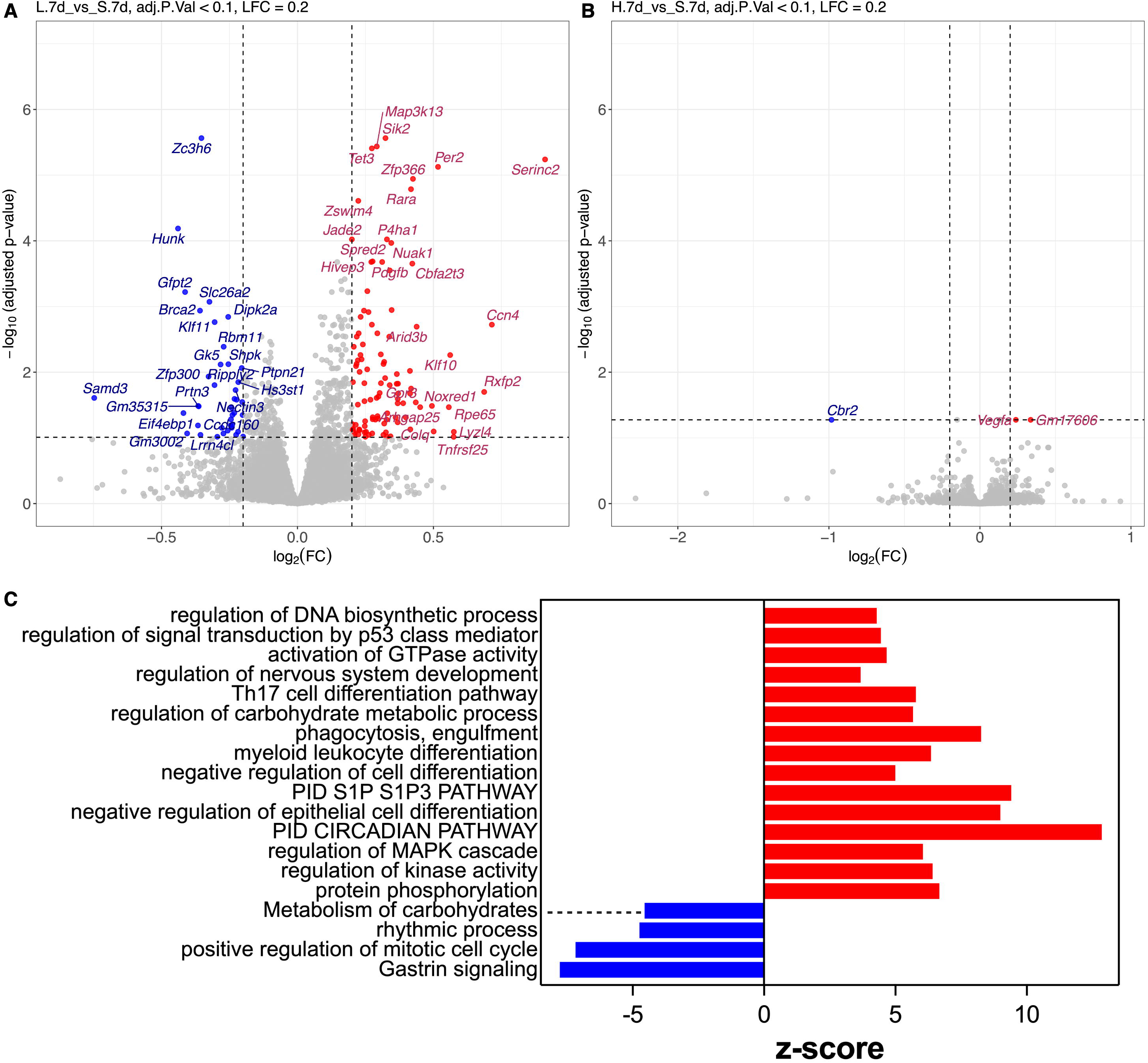
Low dose psilocybin alters transcriptome for 7-days. (A,B) Bulk RNA-seq volcano plots. Downregulated (blue) and upregulated (red) CNS transcriptional changes 7-days following (A) 0.25 mg/kg i.p. psilocybin compared to vehicle (0.9o/o saline) and (B) 1 mg/kg i.p. psilocybin compared to vehicle. Log2 fold-change (FC) denotes magnitude of transcript differential expression on the x-axis. -log10(adjusted p-value) shows FDRq corrected p-values. (C) Gene Ontology (GO) analysis characterizes the most significantly down- and upregulated pathways 2-hours following low dose psilocybin. Z-score denotes Z-standard deviation away from expected counts on the x-axis, and GO pathways on the y-axis.

### Low dose transcriptome alterations outlast high dose

To assess potential lasting changes to the transcriptome following psilocybin treatment, we next assessed the effects of low and high dose psilocybin 7 days after treatment. Combined analysis of both sexes showed a persistence of the transcriptomic response to low dose psilocybin but an attenuation of the response to high dose (Fig 2A-B). While circadian changes continued to be a prominent feature, many of the cellular processes altered by low dose psilocybin at 7 days were distinct from prior timepoints. Notable differences included pathways related to the immune system. Pathways associated with phagocytosis and differentiation of myeloid-lineage leukocytes were upregulated 7 days following low dose psilocybin (Fig 2C). The divergence of the transcriptomic response from prior timepoints, and their absence in high dose treated animals, suggests that low dose psilocybin may have unique molecular effects.

### Females respond more rapidly to low dose psilocybin

Given reports of sex specific differences in dendritic spine formation and elimination following psilocybin administration^16^, we investigated whether a sex dependent effect could be observed in our transcriptomic data. When we performed a sex specific analysis in mice treated with low dose psilocybin, we observed robust changes in female but not male mice at 8 hours, which contrasted with an earlier, combined sex analysis that showed minimal changes (Supp Fig 1A). In females, hundreds of DEG transcripts were both down- and upregulated compared to vehicle treated females at the same timepoint (Fig 3A). In males, there were no changes in differential gene expression compared to vehicle treated males at 8 hours (Fig 3B). Predicted protein interaction networks from the downregulated DEGs following low dose psilocybin in females included proteins involved in cell motility and division, cadherins involved in cell adhesion, and ion channels associated with neuronal membrane potential (Fig 3C). Proteins in the upregulated networks contained numerous receptor subtypes of ionotropic and metabotropic signaling, and proteins involved in chromatin regulation (Fig 3D). Low dose psilocybin altered a diverse set of pathways, with functions spanning metabolic processes, cytoskeletal organization, cell signaling and proliferation (Fig 3E). Cell communication is the global category under which most pathway changes occurred.

**Figure 3.**
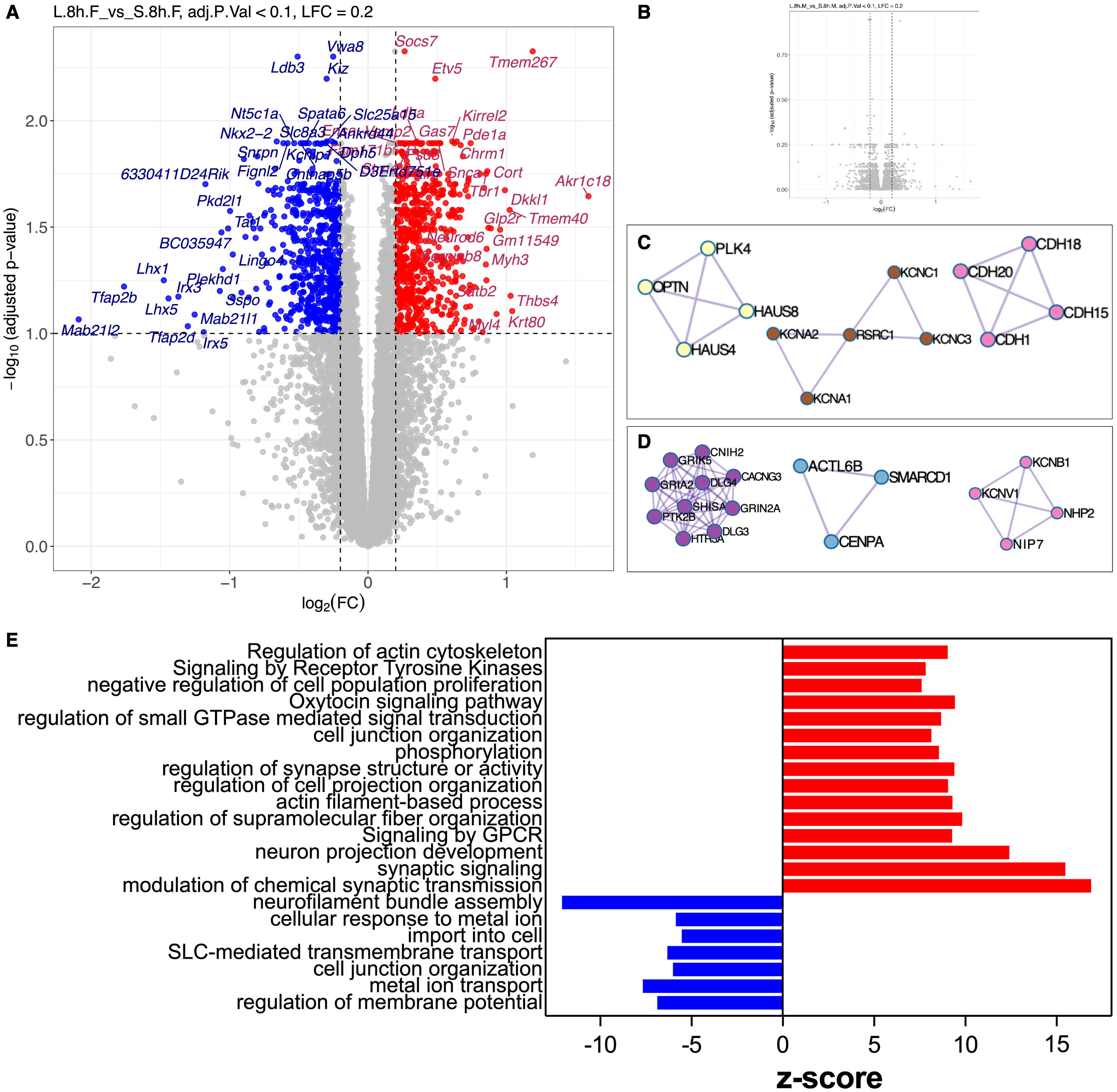
Low dose psilocybin induces robust transcriptional changes in female mice at 8-hours. (A,B) Bulk RNA-seq volcano plot. Downregulated (blue) and upregulated (red) CNS transcriptional changes 8-hours following (A) 0.25 mg/kg i.p. psilocybin compared to vehicle (0.9% saline) in (A) female and (B) male C57BU6J mice. Log2 fold-change (FC) denotes magnitude of transcript differential expression on the x-axis. -log10(adjusted p-value) shows FDRq corrected p-values. (C,D) Predicted protein-protein interactions of (B) downregulated and (C) upregulated DEGs (E) GO analysis characterizes the most significantly downregulated and upregulated pathways 8-hours following low dose psilocybin in females. Z-score denotes Z-standard deviation away from expected counts on the x-axis, and GO pathways on the y-axis.

### Psilocybin induces a sex and dose dependent effect on transcriptome at 8 and 24 hours

We hypothesized that this sex dependent effect would generalize across doses and timepoints. Analysis of both sexes at 8 hours showed that treatment with high dose psilocybin induced dozens of DEGs involved in neurogenesis and nucleosome assembly (Supp Fig 1). However, as with low dose psilocybin (Fig 3), a second analysis of the high dose subset by sex revealed significant differences in the transcripts expressed between sexes and a greater magnitude of differential gene expression in females (Fig 4). The female response to high dose psilocybin evoked hundreds of DEGs at 8 hours (Fig 4A), whereas the males showed a relatively muted response with far fewer down- and upregulated DEGs compared to vehicle (Fig 4B). The overlap in gene expression was minimal, with males and females only sharing 2 upregulated DEGs at 8 hours. Pathway analysis reflected these differences in responses. Females showed significantly greater changes in cell communication and morphogenesis (Fig 4C) whereas males showed changes in metabolism and cell survival (Fig 4D). Given that females responded more robustly to low and high dose psilocybin at 8 hours, it was surprising to see this trend continue for high but not low dose females. We found that the transcriptomic changes seen at 8 hours in low dose females were largely attenuated by 24-hours (Fig 5A) while robust changes were observed in low dose males (Fig 5B). Male and female differential gene expression in response to low dose psilocybin at 24 hours showed only a single overlapping transcript (Supp Fig 3A), however, high dose psilocybin showed significant overlap (Supp Fig 3B). This intriguing sex dependent effect suggests that while males and females respond to high dose psilocybin on a similar time course, their response to low doses is temporally shifted, with females responding and subsequently attenuating their response more rapidly than males. The impact of high dose psilocybin in females at 24 hours continued to show significantly greater magnitude of DEGs compared to males (Fig 5C-D). Pathway analysis of low and high doses at 24 hours further reveals differential responses in females and males (Supp Fig 4).

**Figure 4.**
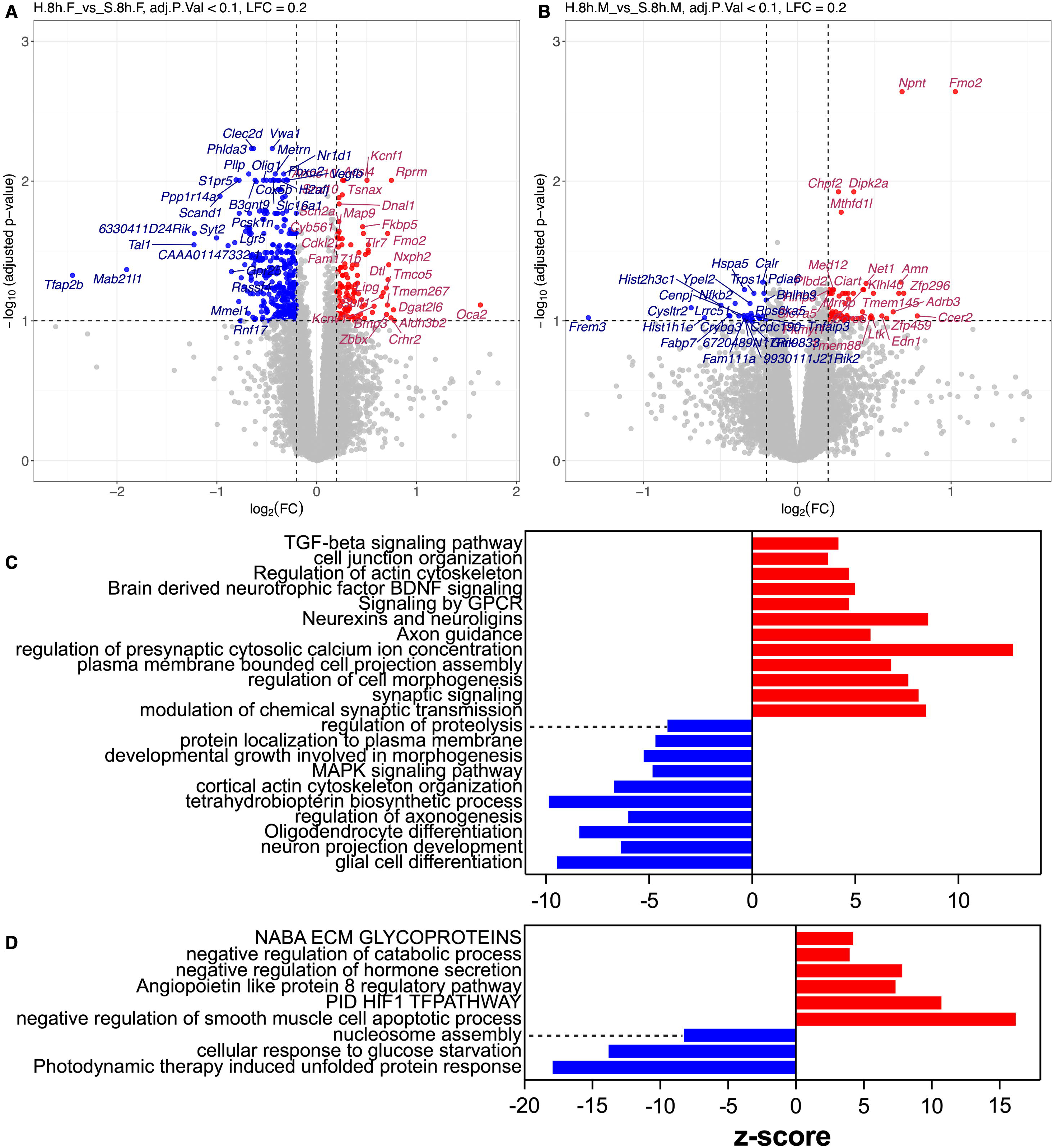
High dose psilocybin induces sex dependent changes at 8-hours. (A,B) Bulk RNA-seq volcano plots. Downregulated (blue) and upregulated (red) CNS transcriptional changes 8-hours following (A) 1 mg/kg i.p. psilocybin compared to vehicle (0.9% saline) in females and (B) 1 mg/kg i.p. psilocybin compared to vehicle in males. Log2 fold-change (FC) denotes magnitude of transcript differential expression on the x-axis. -log10(adjusted p-value) shows FDRq corrected p-values (C,D) GO analysis characterizes the most significantly downregulated and upregulated pathways 8-hours following high dose psilocybin in (C) female and (D) male C57BU6J mice. Z-score denotes Z-standard deviation away from expected counts on the x­ axis, and GO pathways on the y-axis.

**Figure 5.**
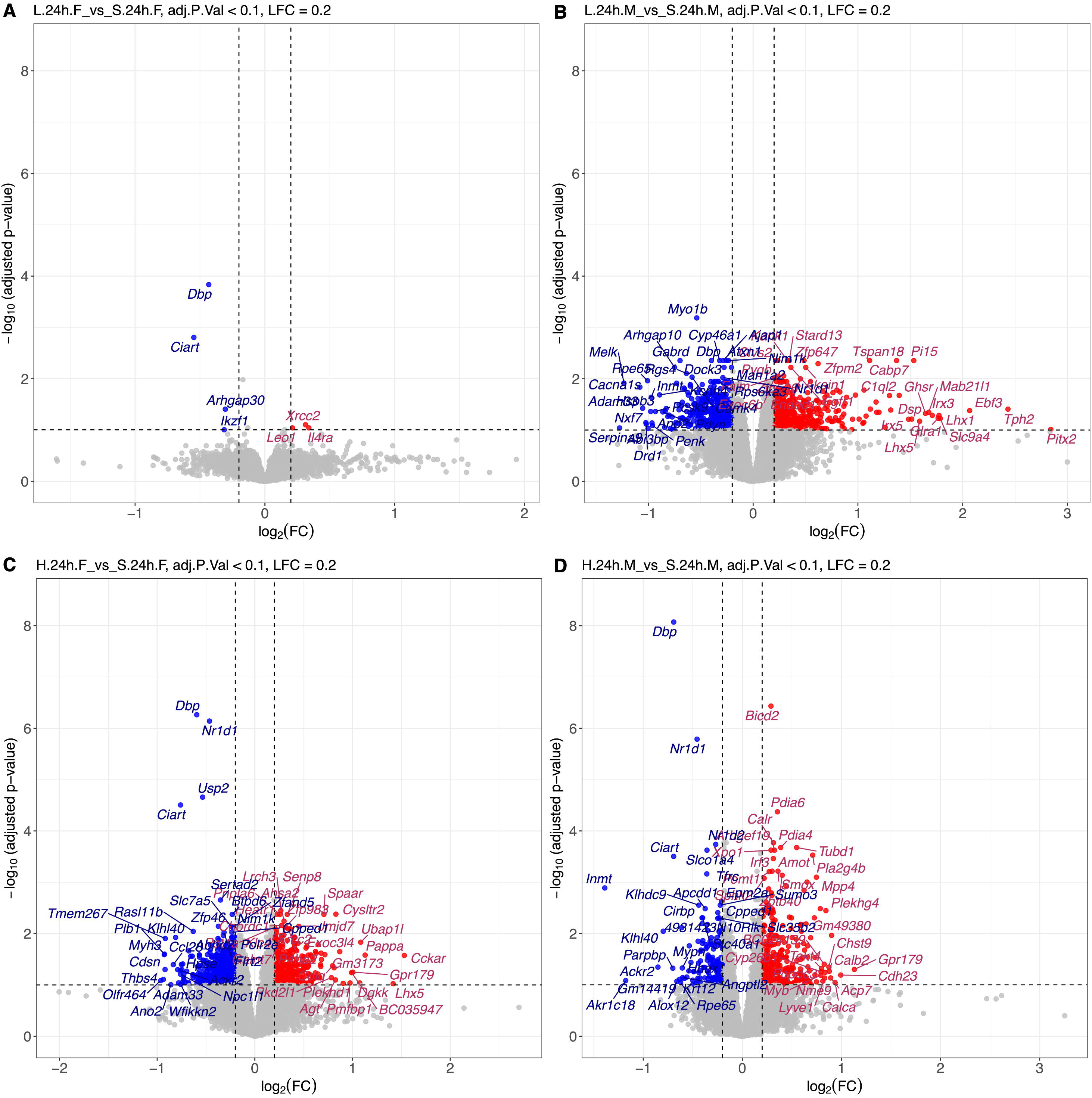
DEGs following low dose psilocybin are attenuated in females but not males at 24-hours. (A,B) Bulk RNA-seq volcano plots. Downregulated (blue) and upregulated (red) CNS transcriptional changes 24-hours following low dose (0.25 mg/kg) i.p. psilocybin compared to vehicle (0.9% saline) in (A) females and (B) males and high dose (1 mg/kg) i.p. psilocybin compared to vehicle (0.9% saline) in (C) females and (D) males. Log2 fold-change (FC) denotes magnitude of transcript differential expression on the x-axis. -log10(adjusted p-value) shows FDRq corrected p-values.

### Effect of low dose psilocybin persistent at 7-days in females and males

To further evaluate effects of low and high dose psilocybin at 7 days, the analysis was subset by sex, revealing that the transcriptomic response to low dose psilocybin persists in both sexes, more in males than females (Fig 6A-B). At 7 days, females only had 18 DEGs (16 up, 2 down) while the males had 135 DEGs (Supp Fig 5C). The transcriptional response had largely attenuated in high dose animals of both sexes (Fig 6C-D). While combined sex analysis of low dose psilocybin demonstrated continued circadian alterations at 7 days (Fig 2), subsequent sex specific analysis revealed that circadian changes at 7 days are driven by male gene expression (Supp Fig 5B). Cell differentiation was present in both sexes, although there were sex-specific enrichments. Both sexes regulated a distinct set of cellular processes compared to prior timepoints, with attenuation of synaptic organization, neuronal differentiation, and ECM remodeling. Distinct pathway activation noted in the combined sex analysis (Fig 2) was driven by gene expression in low dose treated males (Supp Fig 5B). These findings suggest that psilocybin treatment regimens may require sex specific considerations for optimal therapeutic response.

**Figure 6.**
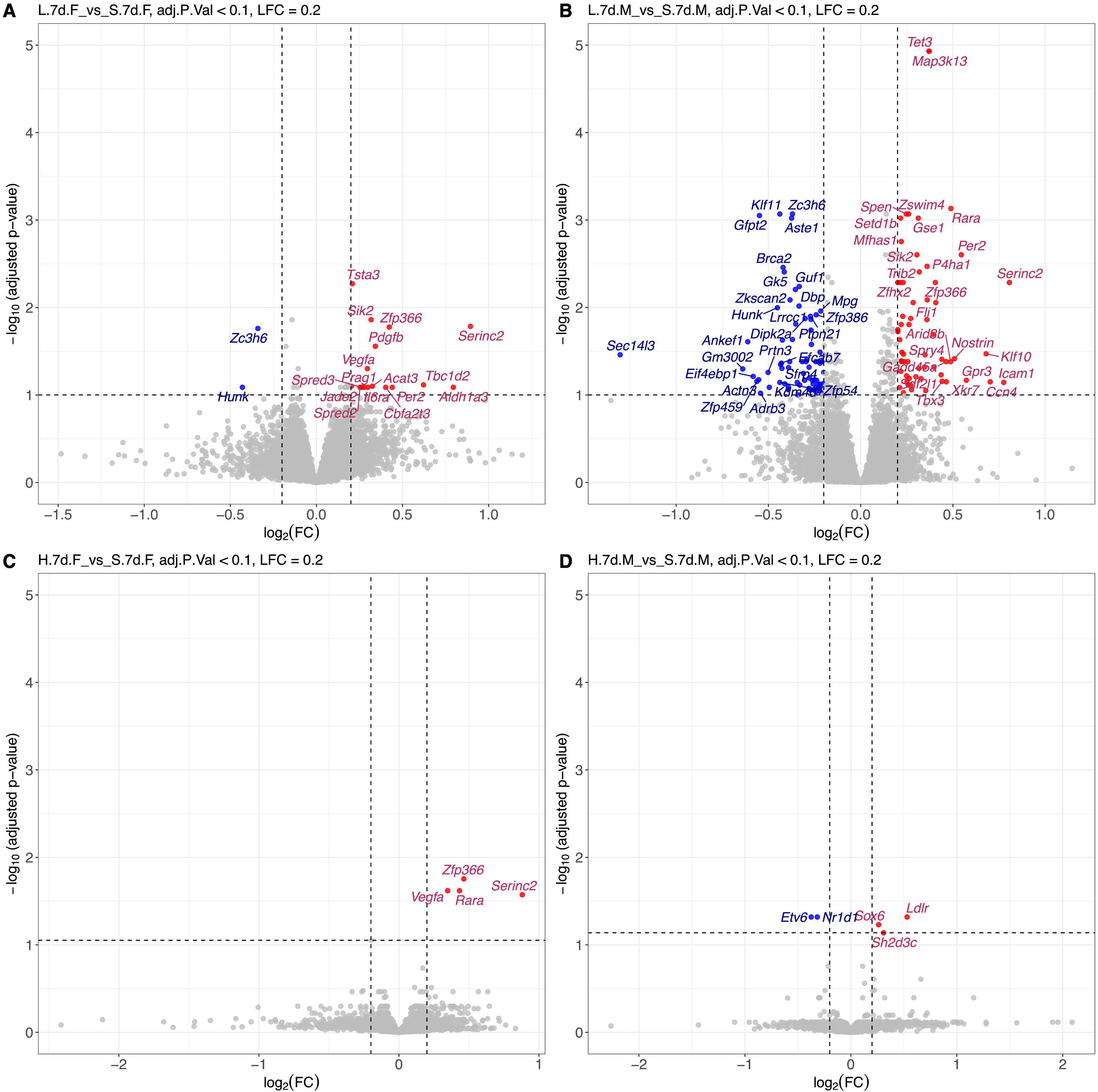
Persistent transcriptomic effect of low dose psilocybin at 7-days. (A,B) Bulk RNA-seq volcano plots. Downregulated (blue) and upregulated (red) CNS transcriptional changes 7-days following low dose (0.25 mg/kg) i.p. psilocybin compared to vehicle (0.9% saline) in (A) females and (B) males, and high dose (1 mg/kg) i.p. psilocybin compared to vehicle (0.9% saline) in (C) females and (D) males. Log2 fold-change (FC) denotes magnitude of transcript differential expression on the x-axis. -log10(adjusted p-value) shows FDRq corrected p-values.

### Weighted gene network correlation analysis reveals psilocybin induced gene hubs

To evaluate expression trends in co-regulated gene groups, weighted gene correlation network analysis (WGCNA) was performed. DEGs from low (0.25 mg/kg) and high (1 mg/kg) dose treated animals from all three timepoints (8 hours, 24 hours, and 7 days) were used to generate dendrograms containing modules of interrelated gene clusters (Supp Fig 6). A signed network approach was used to emphasize positive correlations, and module-trait relationships were explored to uncover biologically relevant pathways and processes. The eigengene is the first principal component of each module and is representative of overall gene expression patterns within each cluster. Female DEGs were clustered into 6 modules of interrelated genes (Supp Fig 6A), while male DEGs resulted in 9 (Supp Fig 6B). The eigengenes of female and male cluster dendrograms were nonoverlapping (Supp Fig 6A-B). Genes with the most connections to other genes within a module are termed hub genes and are considered key regulators of activity within a module. Networks involved in circadian rhythm (Fig 7A) and dendritogenesis (Fig 7B) are illustrated in females, and neuronal and non-neuronal cell differentiation are illustrated for males (Fig 7C-D). Gene ontology analysis of DEGs contained within female and male modules converged on numerous biological processes, including circadian rhythm, neuronal differentiation, MAPK signaling, extracellular matrix remodeling, and modulation of chemical synaptic transmission.

**Figure 7.**
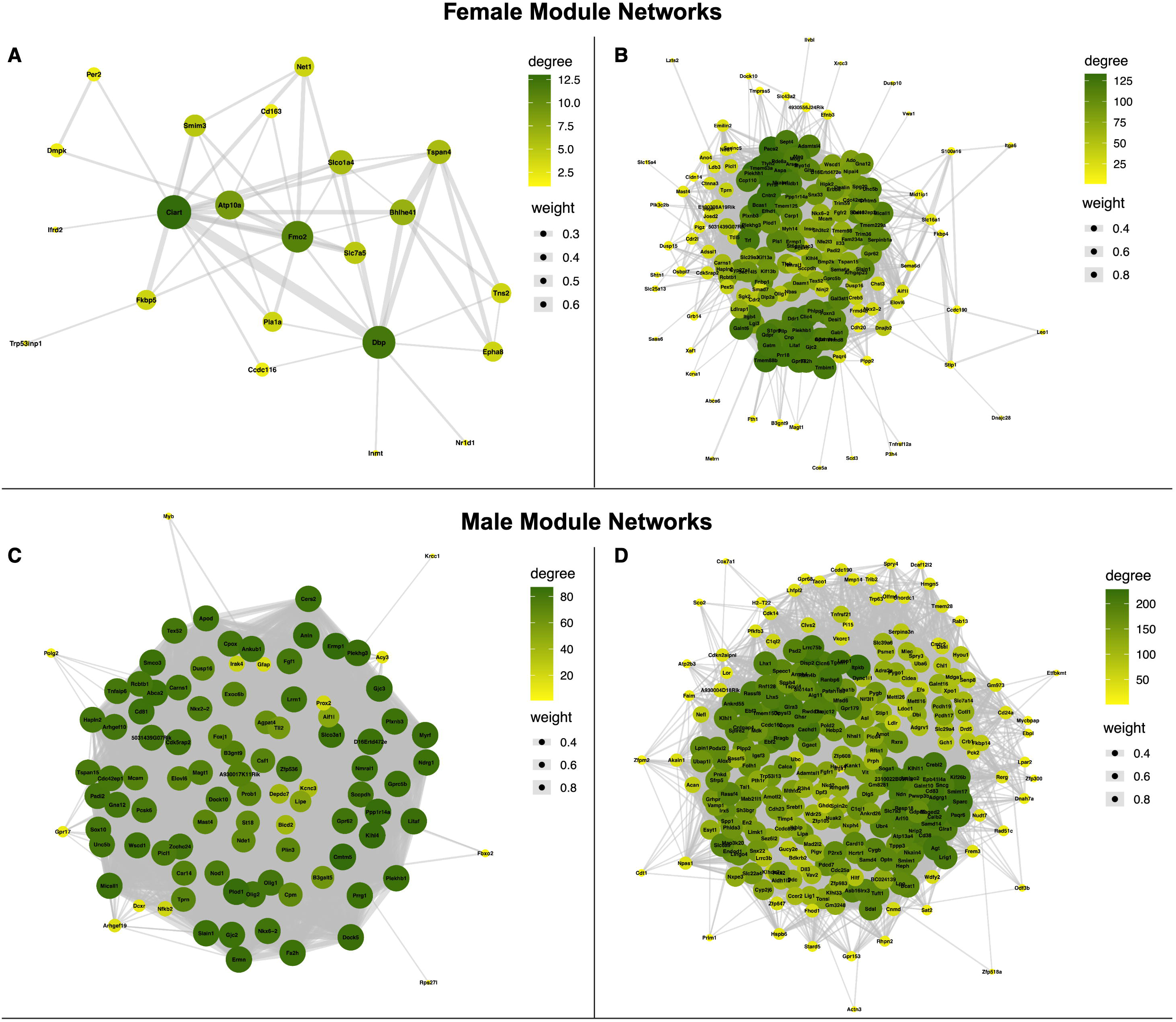
Female and male weighted gene correlation network analysis of psilocybin induced gene expression. Hub gene networks resulting from a signed weighted gene correlation network analysis (WGCNA) subset to all DEGs from the low (0.25 mg/kg) and high (1 mg/kg) dose i.p. psilocybin compared to vehicle (0.9% saline). Adjacency threshold is set to 0.2, and only gene pairs with a correlation higher than 0.2 remain connected in the network. (A-B) Gene networks from the (A) Red and (B) Brown female cluster dendrogram modules. (C-D) Gene networks from the (D) Green and (D) Blue male cluster dendrogram modules. Degree represents the size of a node, which is the number of edges (connections) that a node (gene) has. Larger nodes have more connections (i.e., they are “hub” genes), while smaller nodes have fewer connections. Weight represents the correlation or co­expression strength between two genes. Thicker edges indicate stronger relationships (higher correlation), and thinner edges indicate weaker relationships.

## Discussion

### Dose, sex, and time dependent differences in psilocybin mediated signaling

This study examined the underlying impact of psilocybin therapy on global gene expression patterns at three timepoints (8 hours, 24 hours, and 7 days) following a single administration of psilocybin. The primary findings are that psilocybin produces distinct and lasting sex- and dose-dependent transcriptional responses in the mouse forebrain. Previous work using a targeted qPCR panel identified plasticity-related transcriptional changes in the male rat prefrontal cortex and hippocampus^31^. Transcriptional changes in the murine male prefrontal cortex have been reported at 24-hours^32,33^, however, the durability, dose, and sex dependence of these genomic changes have not. In the present study, analysis of low (0.25 mg/kg) and high (1 mg/kg) dose psilocybin revealed differential patterns of gene expression over time and suggest that differences in dose could greatly influence the mechanism of psilocybin’s therapeutic action. The high dose resulted in expression of a broad set of DEGs, impacting pathways associated with neurogenesis, nucleosome assembly, circadian rhythm, and neuronal and glial cell differentiation. The low dose resulted in fewer but sustained transcriptional changes, including immune cell differentiation and alterations to circadian genes out to 7 days. These findings indicate that dosing of psilocybin cannot be thought of as a “volume knob”, with lower doses producing dampened effects of higher doses. Different doses of psilocybin produce distinct effects that may uniquely alter neuronal and non-neuronal signaling. The relevance of these distinct biological effects to therapeutic efficacy is unclear and warrants further investigation.

This study also revealed pronounced sex specific differences in differential gene expression, with females producing and subsequently attenuating a transcriptomic response to low doses more rapidly than males. Females also responded more robustly to high doses than males at 8 and 24 hours, with signal attenuation in both sexes by 7 days. Our findings provide additional mechanistic insight to reports of sex specific differences in behavior following psilocybin exposure^34,35^. Together, these data indicate that optimal therapeutic dosing likely varies by sex, raising important questions for the design of clinical trials. The earlier and more robust female response suggests a potentially greater therapeutic effect for certain psychiatric conditions. Hormonal or genetic differences influencing receptor density, distribution, or intracellular signaling pathways may be the basis of the sex specific differences reported here. Future studies may reveal the molecular mechanisms that underly these sex specific differences– as well as their significance for personalized approaches to psilocybin therapy.

### Implications for future clinical trials

There is considerable interest in the use of daily, low dose (i.e. subperceptual), psilocybin to produce antidepressant effects. Clinical data on lower doses is limited and demonstrates modest effects on mood, confounded by expectation bias^36^. Relatively little is known about the mechanistic differences between different doses of psilocybin, or if therapeutic benefit could be achieved with a lower dose. In this study, a notable and unexpected observation was the persistent transcriptional effect of low-dose psilocybin at 7 days, which outlasted high-dose changes and suggests that dosing paradigm impacts the duration of psilocybin’s biological effects. A similar dose-dependent effect on gene expression has been observed for ketamine, where the effects of low doses persist beyond the high dose^37,38^. This phenomenon could be the result of receptor downregulation in response to high doses, whereas low doses have no such effect. Dose dependent differences in receptor class engagement is a well-established paradigm in pharmacology, and the effects of lower doses of psilocybin may be mediated by a combination of receptors, including the TrkB, 5-HT_2A_ and other 5-HT receptor subtypes. The therapeutic usefulness of persistent molecular changes following low dose psilocybin is unknown. These changes may be the basis for anecdotal reports on the efficacy of low dose psilocybin for mood enhancement.

Low dose regimens are also of interest in the clinical trial setting where large doses often induce unwanted perceptual effects, headache, and nausea^39^. As with any pharmacologic agent, the balance of wanted and unwanted effects can be managed by determining the lowest dose capable of producing a desired therapeutic outcome. These data provide clarity on the molecular effects of lower doses of psilocybin and may be informative to researchers querying whether a subperceptual dosing paradigm is a viable approach to achieve an antidepressant and/or a neuroplastic effect. The molecular overlap of low and high doses reported here suggests that lower doses of psilocybin may possess some of the therapeutic properties of higher doses. While the significance of these molecular changes to clinical outcomes remains unknown, their overlap with the efficacious higher doses should encourage future clinical trials to examine whether lower doses of psilocybin produce antidepressant effects in humans.

Given a growing body of literature implicating inflammation and immune dysfunction in depression^40–43^, immunomodulatory effects could prove useful in treating inflammatory subtypes of depression or non-psychiatric inflammatory conditions. The upregulation of pathways associated with differentiation of myeloid-lineage leukocytes and phagocytosis 7 days following treatment should encourage further investigation into a possible immunomodulatory effect of low dose psilocybin. As psychiatric disorders are commonly disrupt the circadian clock^44^, our observation of altered circadian genes at 8 and 24 hours is notable and merits further investigation. *Ciart* is a circadian associated repressor of transcription, and its downregulation promotes expression of *Bmal1* and *Clock*, facilitating healthier sleep architecture by reducing fragmented sleep and increasing depth^45–47^.

In conclusion, these data revealed sex dependent molecular effects of psilocybin and support previous studies demonstrating its effect on dendritogenesis. Differential gene expression impacted a diverse set of biological processes, but common features included functions related to neuronal differentiation, neurogenesis, and changes in receptor signaling. Our report of a sex- and dose-dependent duration and magnitude of effect will be useful for the optimization of treatment paradigms for PTSD, TRD, and other psychiatric conditions. This dataset provides fundamental insights to the diverse molecular effects of psilocybin and can be queried using a web-based tool developed by the Fryer lab: https://fryerlab.shinyapps.io/psi1/

### Limitations

This study employed bulk RNA sequencing (RNA-seq) of forebrain samples to assess the murine response to psilocybin, which presents several limitations. Bulk RNA-seq data reports the averaged gene expression across all cell types within a sampled tissue, which obscures cell specific patterns of gene expression. The use of forebrain presents two limitations: first, it excludes brain stem and cerebellar components and biases the analysis toward forebrain structures; second, it limits spatial resolution by assessing the transcriptome of a broad anatomical region containing heterogenous neuronal and non-neuronal cell populations. Murine models do not fully model human drug responses, given pharmacokinetic and pharmacodynamic differences. Future studies with single-cell/nucleus or spatial transcriptomics could address some of these limitations and further advance our understanding of psilocybin’s action in the brain.

## Data availability

Data are available within the manuscript body and supplementary materials. Upon manuscript acceptance, code used to analyze bulk RNAseq and raw fastq files will be available via GEO and SRA.

## Acknowledgements

We would like to thank the National Institute on Drug Abuse, Drug Supply Program (Bethesda, MD, USA) and the Mayo Clinic Rochester Genome Analysis Core.

## Author contributions

J.H. Barnett and J.D. Fryer conceptualized the study and its methods, performed the experiments, and collected the data. J.H. Barnett and K.T. Todd were responsible for data curation. K.T. Todd performed the formal analysis. J.D. Fryer secured funding and supervised all aspects of the project. J.H. Barnett was responsible for project administration, animal procurement and care, and tissue analysis. J.H. Barnett was responsbible for data visualization and figure assembly. J.H. Barnett, J. Benetatos, and J.D. Fryer contributed to the interpretation of experiments. J.H. Barnett wrote the original draft of the manuscript. J.H. Barnett, J. Benetatos, K.T. Todd, B.E. Rabichow, K.A. Gibson, K.C. Olney, and J.D. Fryer edited and revised the manuscript.

