## Supplementary figures and images for "Psilocybin causes sex, time, and dose dependent alterations in brain signaling pathways"

### Supp Fig 1

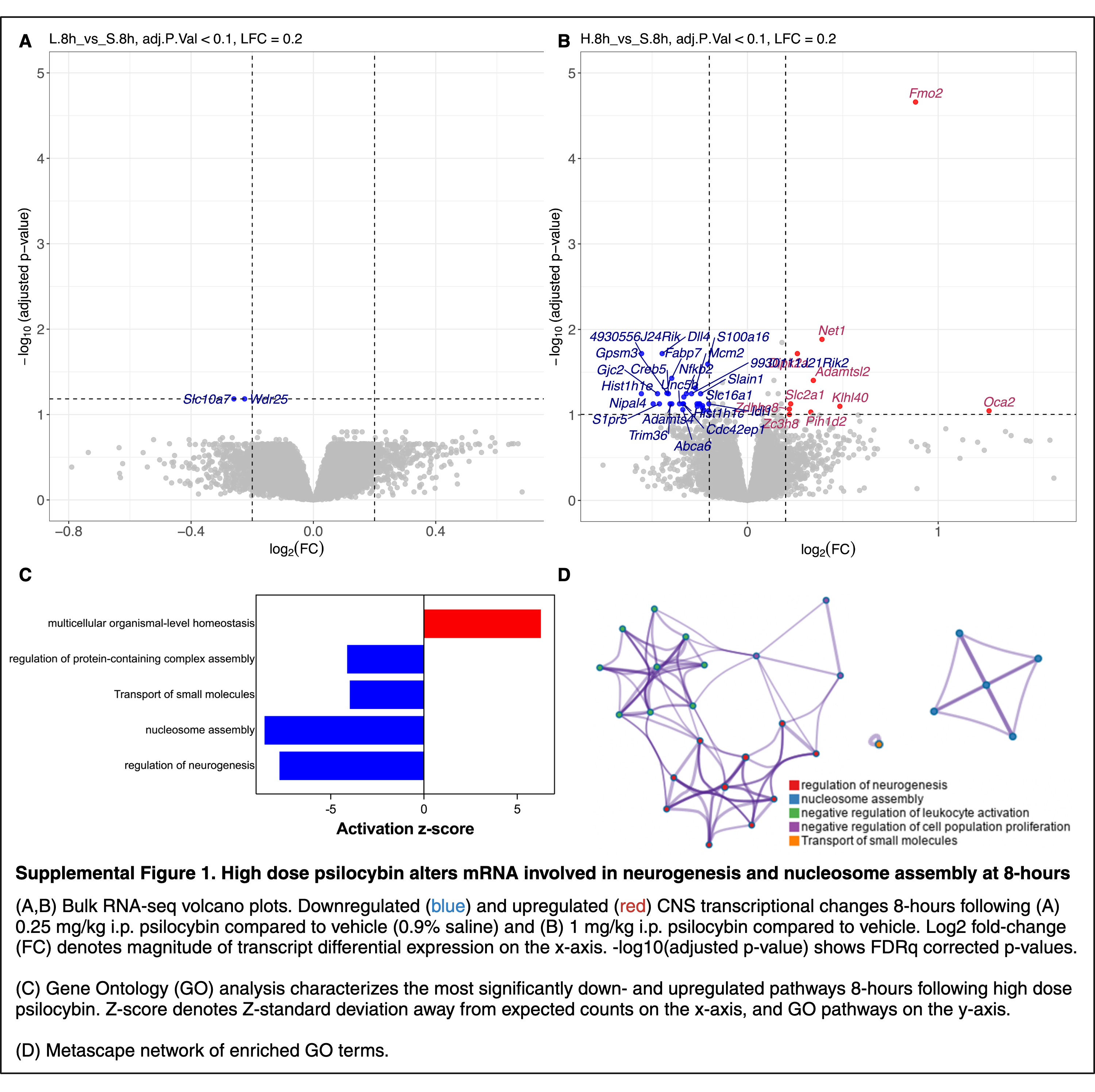

### Supp Fig 2

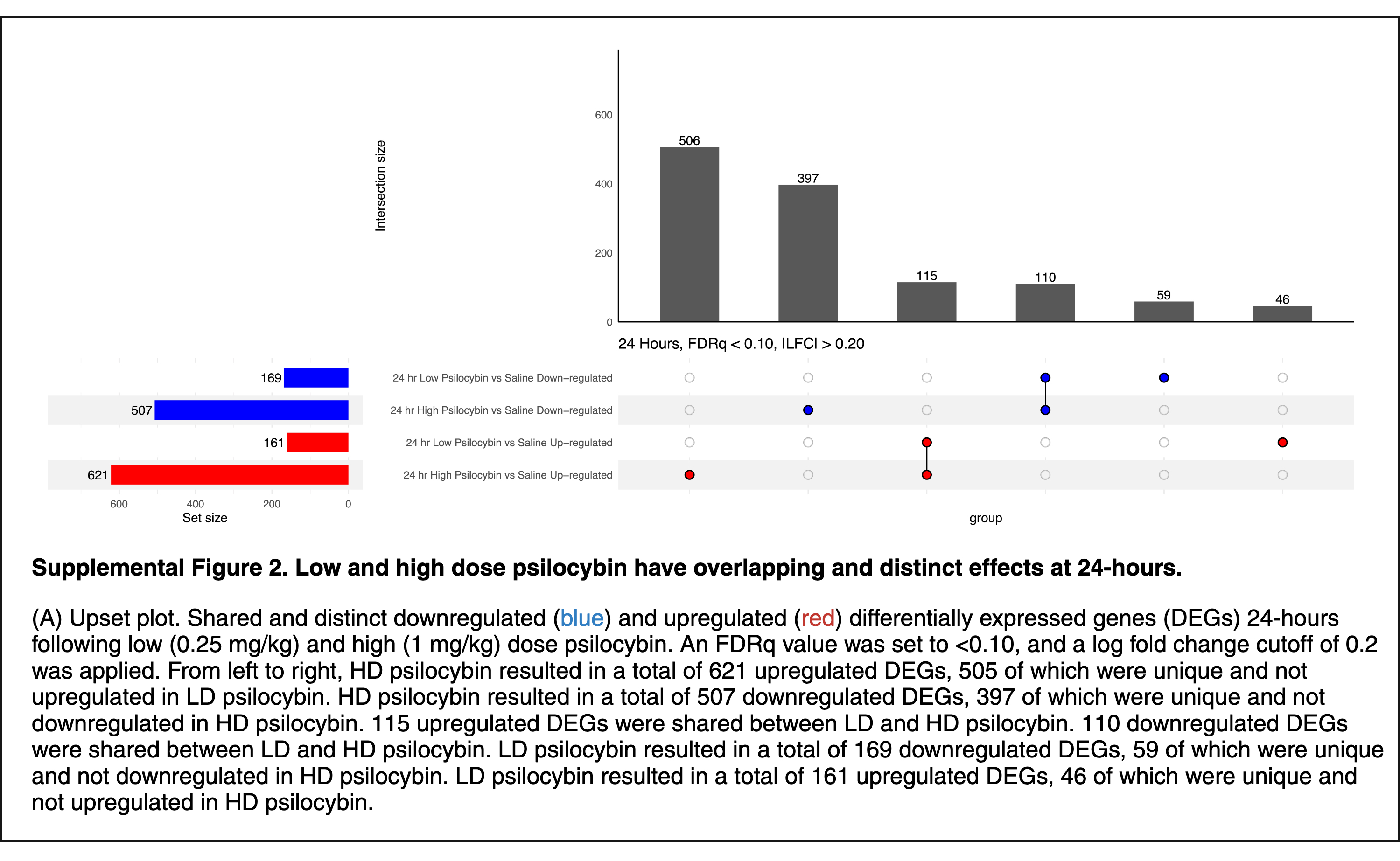

### Supp Fig 3

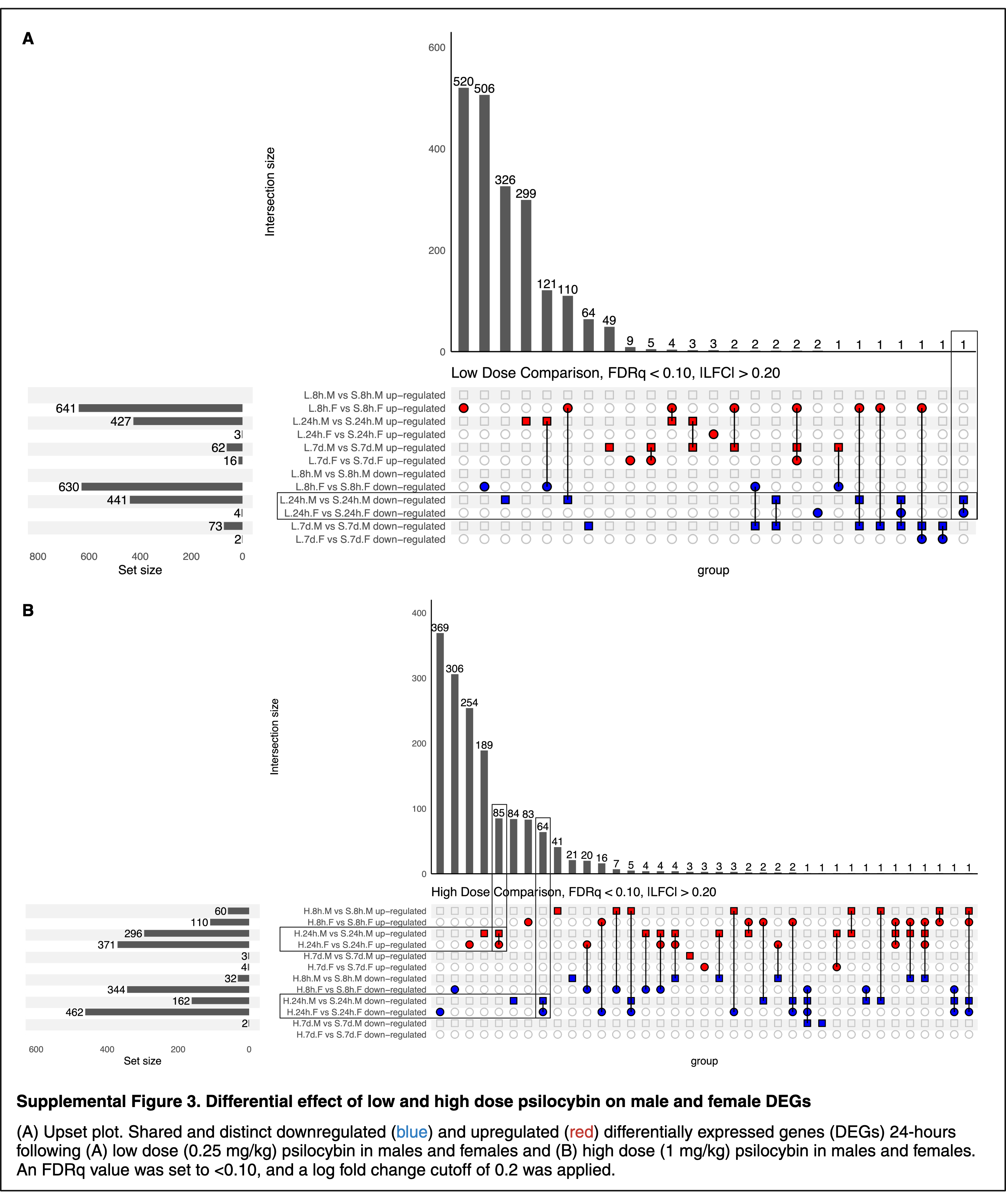

### Supp Fig 4

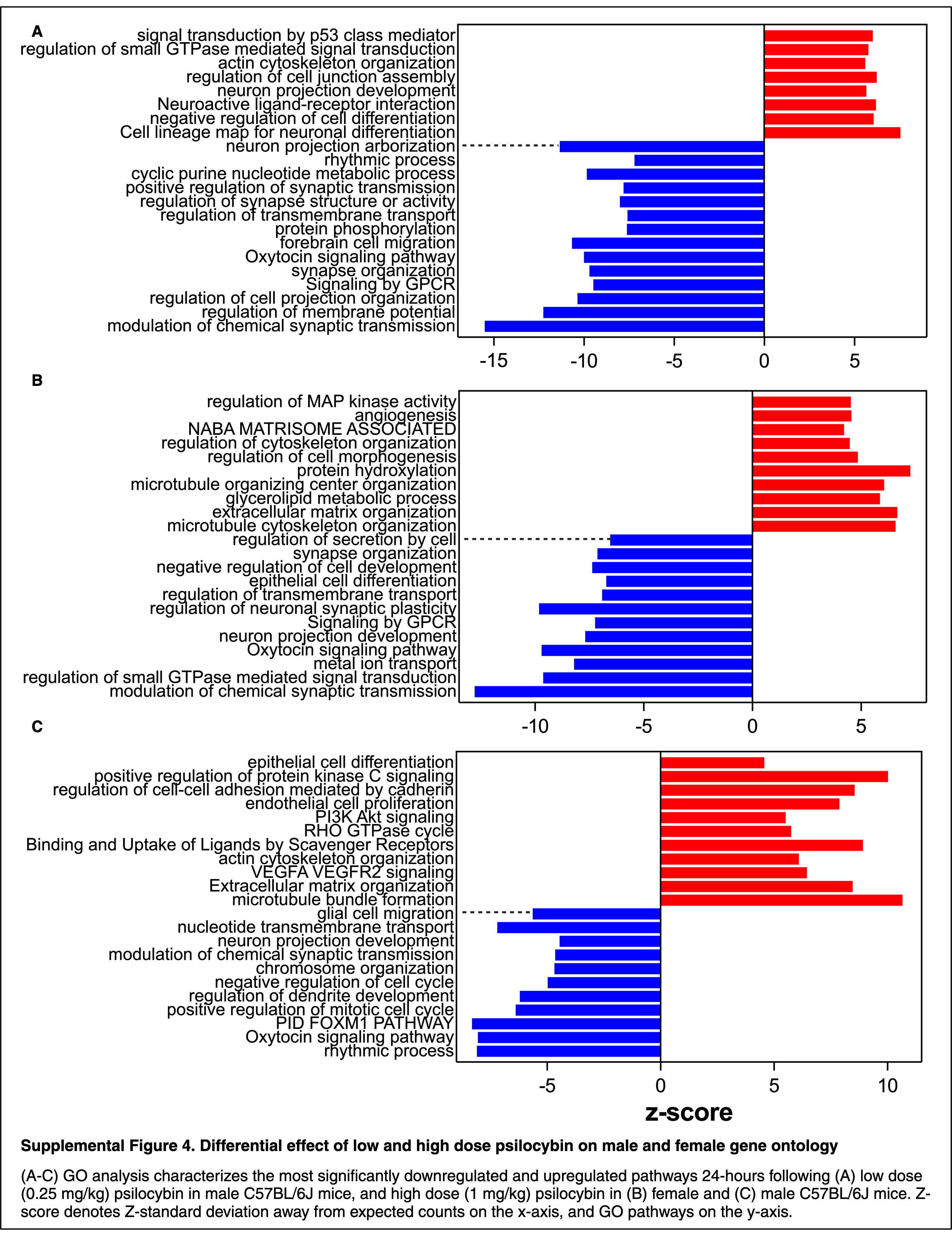

### Supp Fig 5

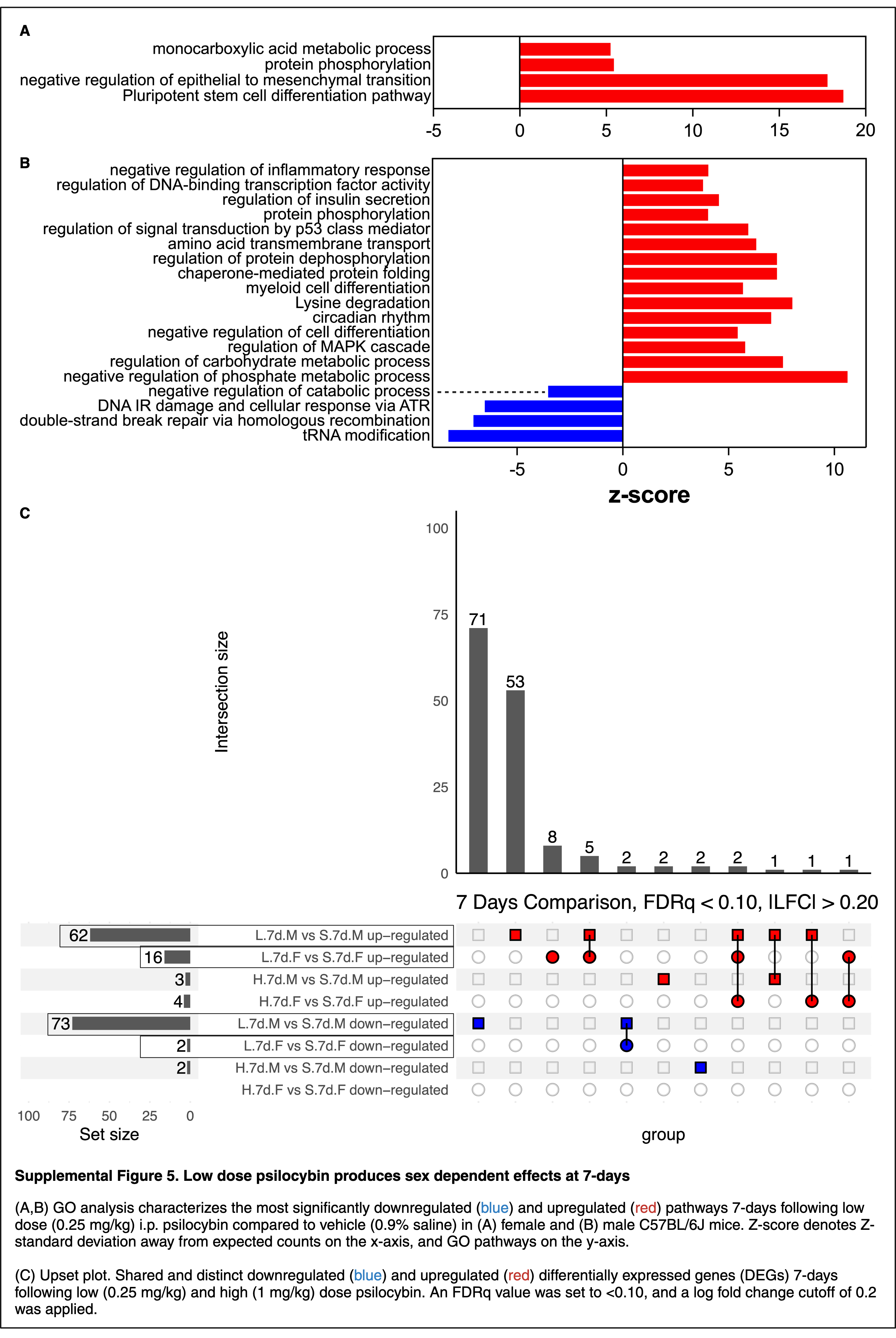

### Supp Fig 6

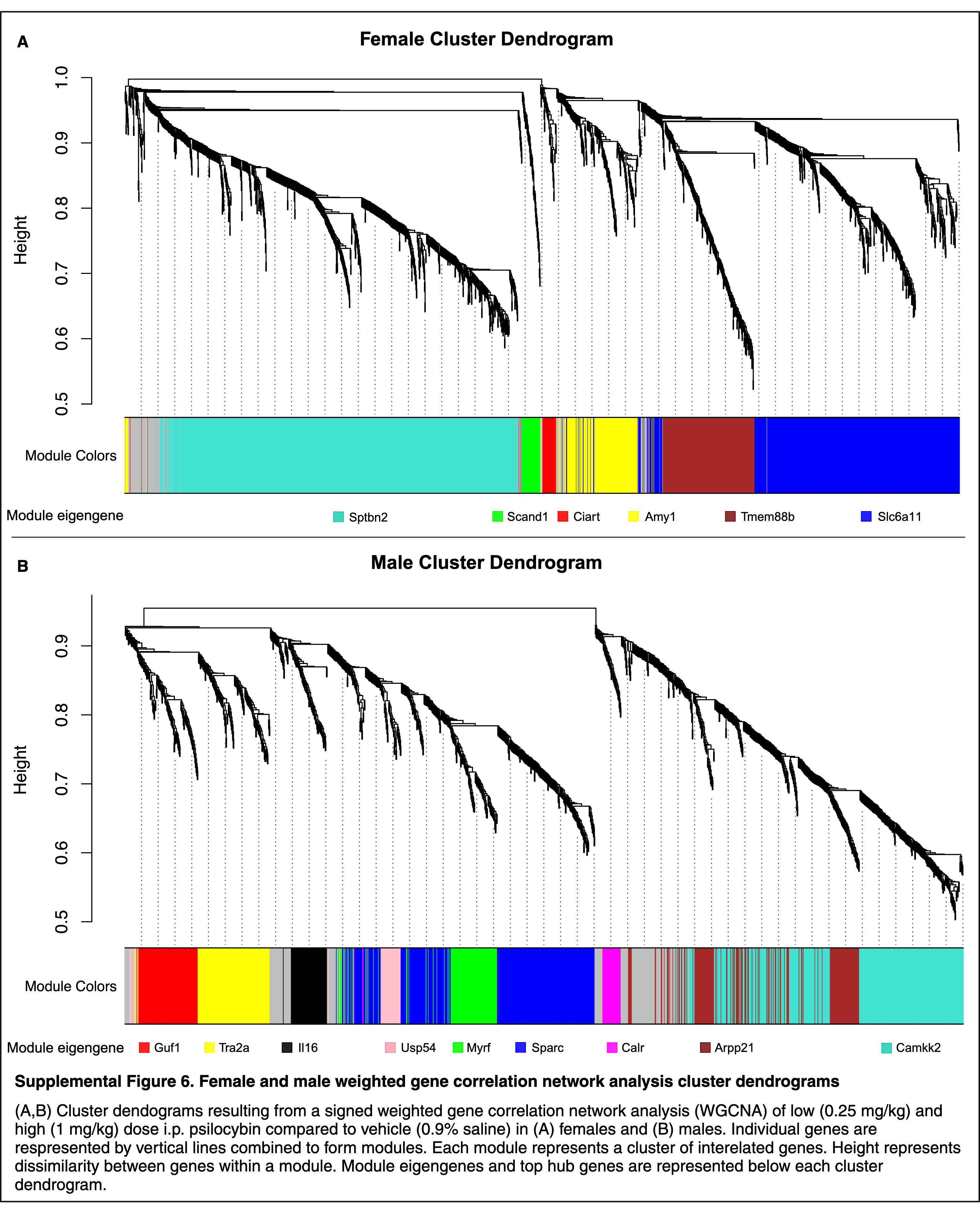
